# Inactivation of *B. cereus* Spores in Whole Milk and Almond Milk by Novel Serpentine Path Coiled Tube UV-C System

**DOI:** 10.1101/2022.02.22.481446

**Authors:** Brahmaiah Pendyala, Ankit Patras, Vybhav Vipul Sudhir Gopisetty, Pranav Vashisht, Ramasamy Ravi

**Affiliations:** Food Biosciences and Technology Program, Department of Agricultural and Environmental Sciences, Tennessee State University, Nashville, 37209, TN, USA

**Author notes:** Co-Corresponding authors, **Ankit Patras, Ph.D**. Associate Professor, Food Science & Engineering, **Brahmaiah Pendyala, Ph.D**. Research Scientist.

**Keywords:** UV-C irradiation, Coiled tube UV system, Whole milk, Almond milk, *B. cereus* spores, Lipid peroxidation, Volatiles

## Abstract

A novel continuous thin-film (1.59 mm) serpentine path coiled tube (SPCT) UV system operating at 254 nm wavelength was designed and compared with flow field distribution of whole milk with helical path coiled tube (HPCT) UV system using computational fluid dynamics. The results revealed efficient velocity magnitude distribution at serpentine bend geometric locations of the SPCT UV system. Further in this study, we evaluated *B. cereus* Spores inactivation in whole milk (WM) and almond milk (AM) using the developed SPCT UV system. Experimental data showed that > 4 log reduction of spores was achieved after six and ten passes of WM and AM at a flow rate of 70 and 162 mL/min, respectively. The biodosimetry method was used to verify the delivered reduction equivalent fluence (REF) and reported as 33 ± 0.73 and 36.5 ± 1.9 mJ/cm2. We noticed no significant effect on lipid oxidation and volatiles profile (p > 0.05) up to delivered REF of 60 mJ/cm^2^. This study demonstrated that high levels of inactivation of *B. cereus* spores could be feasible with minimal impact on product quality by UV-C processing of dairy and non-dairy opaque scattering fluids.

## 1. Introduction

Psychrotrophic spore-forming bacteria (E.g., *Bacillus cereus*) that can grow at temperatures < 7 °C is a significant food safety and spoilage concern for dairy and plant-based low acid (pH > 4.2) milk products (E.g., WM and AM) (Georget et al., 2014, Gopal et al., 2015). Spores of these bacteria can survive and germinate at pasteurization and refrigeration temperatures, causing safety and spoilage issues (Griffiths, 1992). Germinated *B. cereus* produces potential pathogenic substances such as; emetic and enterotoxins, hemolysins, metalloproteinases, phospholipase C, collagenases, and beta-lactamases (Turnbull et al., 2002). *B. cereus* was responsible for 63,400 foodborne illnesses in the United States between 2000 and 2008 (Scallan et al., 2011) and posed a health risk. World health organization (WHO) estimated that the *B. cereus* caused foodborne illnesses of 256,775 (95 % UI 43,875–807,547) in 2010 (Kirk et al., 2015). On the other hand, *B. cereus* cause off-flavors such as bitty cream and sweet curdling in milk due to its enzymatic activities, contributing to the dairy industry’s financial losses by product spoilage (Gopal et al., 2015).

Ultra-high temperature (UHT) processing or ultra-pasteurization (138 °C for 2 seconds) (https://www.idfa.org/pasteurization) is a traditional technique used to inactivate thermophilic bacterial endospores in low acid liquid foods (E.g., milk). However, exposure to high-temperature results in the deterioration of nutritional (reduction of bioactives, loss of vitamins, protein denaturation, enzyme inactivation, and lipid oxidation) and sensory (color, texture, and flavor) quality of milk (Cappozzo et al. 2015; Popov-Raljić et al. 2008; Renner, 1988; Van Boekel, 1998). In addition, UHT is a high energy-consuming process and influences the product’s final value to ensure economic sustainability (Delorme et al. 2020). Non-thermal sterilization techniques offer promising UHT alternatives for processing dairy and plant-based milk products with enhanced quality. Numerous research studies have been conducted to evaluate non-thermal sterilization methods; high-pressure processing, ultraviolet (UV-C) and pulsed light, pulsed electric field, ultrasonication, and cold plasma (Li & Farid, 2016).

Among these, UV-C light processing has been considered one of the most promising technologies due to its low energy consumption, and the system can be easily retrofitted with aseptic packaging. In addition, it does not generate any chemical residues or toxic chemical byproducts (Patras et al., 2021). In 2016, the European Union commission approved UV-C treated (1045 J/L) milk to obtain extended shelf-life (EFSA NDA Panel, 2016). In contrast, the FDA has not approved UV-C light usage for milk treatment, and it is also regulated through the PMO. Perhaps more robust UV studies are required to understand UV technology from food safety and quality perspectives. Antimicrobial properties of UV-C light at 254 nm have been extensively studied against vegetative bacteria, spore forms, viruses, fungi, algae, and protozoa (Malayeri et al., 2016). UV-C germicidal light has a great potential to inactivate various bacterial spores (Mamane-Gravetz and Linden, 2004, Blatchley et al. 2005, Choudhary et al., 2011, Bandla et al., 2012, Gayán et al., 2013, Pendyala et al., 2019 & 2020, Ansari et al., 2019). UV-C treatment of spores produces spore photoproducts (SP) as a significant product with low CPD and 64PP, resulting in DNA damage (Moeller et al., 2007, Setlow, 2014).

Low UV light penetration within high UV absorbing opaque liquids could result in inefficient microbial inactivation using systems with high optical depth (Pendyala et al., 2021). In order to increase light penetration, researchers have developed and engineered different UV systems. These UV systems involve either thin-film or ultra-thin-film reactors with film thickness ranges from 0.02 to 5 mm (Pendyala et al., 2021). The UV systems were primarily designed to reduce the optical thickness of fluids, thereby improving their treatment efficacy (Koutchma et al., 2004, Ye et al., 2008, Crook et al. 2015, Martinez-Garcia et al., 2019, Pendyala et al., 2021). Some researchers designed and developed coiled tube Dean flow reactors to create Dean forces and turbulence at low Reynolds number (< 2100) and thereby dose distribution (Dean, 1927, Choudhary et al., 2011, Bandla et al., 2012, Gopisetty et al., 2019, Ansari et al., 2019, Pendyala et al., 2020, Vashisht et al., 2021). UV inactivation of spores in skimmed cow milk, raw milk, and soymilk have been described using HPCT continuous flow UV units (Choudhary et al., 2011, Bandla et al., 2012). However, these studies did not evaluate the absorbed UV fluence and reported surface dose at experimental conditions. Similarly, Ansari et al. (2019) investigated UV-C inactivation of *B. subtilis* spores in skim cow milk and estimated absorbed UV fluence using potassium iodide/iodate-based chemical actinometry method. However, since the optical properties of milk differ from potassium iodide/iodate, the absorbed fluence would differ from reported values. Test fluid optical attenuation properties (absorbance, scattering, reflection) and lamp UV intensity are essential parameters to consider while designing the UV system’s optical path length (Patras et al., 2021). In addition, fluid viscosity, density, and coil diameter that can significantly influence Reynolds number and Dean number are crucial parameters to be considered for the design and development of efficient mixing UV systems (Patras et al., 2021).

Computational fluid dynamics (CFD) application has been widely used in the design of UV reactors (Sozzi and Taghipour, 2006, Atilgan, 2013, Younis et al., 2018, Mandal and Pratap-Singh). CFD predicts the fluid flow based on the laws of conservation of mass, momentum, and energy and considers reactor geometry, physical properties of the fluid, and boundary conditions of a flow field (Scott and Richardson, 1997). We hypothesized that using the thin-film SPCT UV system can significantly increase light penetration and fluence distribution due to strong dean vortices at serpentine bends. Also, considering the optical properties of test fluid for evaluating fluence delivery by biodosimetry would be a suitable method for system validation studies. The major objectives of this study are (i) comparison of fluid flow in SPCT and HPCT UV reactors using CFD; (ii) validate SPCT reactor’s efficacy to inactivate *B. cereus* spores in WM and AM; (iii) evaluate the quality of the UV-C processed milk via lipid peroxidation volatiles analysis.

## 2. Materials and Methods

### 2.1 Microbial strains and culture conditions

*B. cereus* ATCC 14579 strain was purchased from the American Type Culture Collection (ATCC) and used in the present study. *B. cereus* ATCC 14579 was propagated using Brain Heart Infusion broth (BHI, Beckton Dickinson, Franklin Lakes, NJ), sporulated using Mineral Salts Medium (MSM), harvested and purified endospores as described in our previous studies (Pendyala et al., 2019, 2020 & 2021).

### 2.2 Preparation of microbial suspensions of WM and AM

The ultra-pasteurized WM was obtained from a local grocery store in Nashville, Tennessee, and AM was obtained from Beber Almond mill, Chico, CA. The purified spores were inoculated at a final concentration of > 10^7^ CFU/mL to prepare the WM and AM test fluids. Before inoculation, background spore populations of WM and AM were examined.

### 2.3 Measurement of optical properties

Optical properties of the WM and AM microbial suspensions were measured using a double beam Cary 100 Spectrophotometer (Varian, USA) equipped with a 6-inch single Integrating Sphere (Labsphere, DRA-CA-30, USA) to estimate scattered light at 254 nm wavelength (Shenoy and Pal, 2008). Thin quartz cuvettes (0.08 mm path-length) were prepared and used to measure optical properties. The transmittance and reflectance (diffuse reflectance) of light were collected by the integrating sphere when the sample was placed at the entrance and exit ports, respectively (Gunter-ward et al., 2018). The amount of light transmitted and reflected by the quartz cuvettes was also quantified and considered to estimate absorption, scattering coefficients, and reflectance. The inverse adding-doubling (IAD) program was used to compute absorption and scattering coefficients by applying total transmittance, reflectance, and refractive index as input values (Prahl et al., 1999). Ultraviolet transmittance (UVT- %/cm), which indicates the fraction of incident light transmitted through a material over a 1 cm path, was calculated as per equation 1.

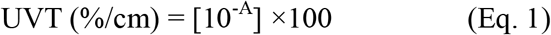

where A represents the absorbance (base_10_) of the test fluid at 254 nm for a 1cm path.

### 2.4 Estimation of delivered fluence by biodosimetry

Estimating UV fluence delivery at experimental flow conditions was performed using the biodosimetry method (Bhullar et al., 2018, Patras et al., 2021, Pendyala et al., 2020, Vashisht et al., 2021). Briefly, the dose-response equations for the WM and AM spore suspensions were determined using dose-response curves obtained from the standard bench-scale collimated beam apparatus. Optical path length was 6 mm and treated the WM, and AM spore suspensions in 10 mL beaker at average UV dose ranges from 0 - 40 mJ/cm^2^ (n = 3). Since WM and AM scatter UV-C light, in addition to test fluid scattering factor (*S*) was estimated by using radiative transport equation (Prahl et al., 1999) and considered for calculation of average UV fluence rate (*Eavg*) and thereby delivered UV dose (*D*) in the stirred sample.

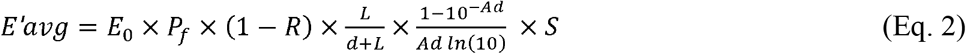

*E*_0_ is the radiometer meter incident irradiance reading at the center top surface of the water in the beaker; *P_f_* is petri factor; 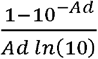 is water factor (1 − *R*) is reflectance factor;, and 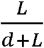 is light divergence factor

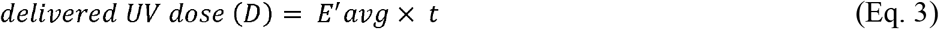

*t* is the exposure time in sec.

Scattering factor (*S*) was estimated. Log-inactivation values from the SPCT reactor were used in the dose-response equation (Eq. 4) to estimate the Reduction Equivalent Fluence (*REF*) (USEPA, 2006).

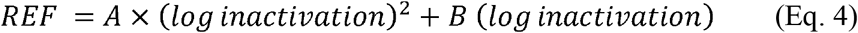

Where *A* and *B* are coefficients obtained from collimated beam system dose-response curve quadratic equation.

### 2.5 UV irradiation treatment

WM or AM microbial suspensions were irradiated using an SPCT UV system (Figure 1). The system consists of a low-pressure mercury UV lamp (15 W) positioned in the middle of the reactor, a cooling fan, a flow tube (Teflon with ≈ 60 % UV-C_254nm_ transmittance) assembly, and a peristaltic pump (Watson-Marlow). The UV lamp was connected to a ballast (power supply), with a cooling fan supplying cool air to maintain the lamp temperature. The Teflon flow tube (1.59 mm diameter) with a length of ≈ 4 meters was assembled by wrapping the tubing through the holes of the welded metal strip onto the reactor body. There were 17 sharp serpentine bends (bend diameter 1 cm) to induce radial mixing. Fluid flow in the serpentine path at sharp bends induces strong dean vortices where the fluid particles are mixed in a clockwise and anticlockwise directions around the bends. Using an Ocean optics sensor, the tube surface UV-C irradiance intensity at a 4.5 mm distance from lamp was measured as 14.85 mW/cm^2^. WM or AM was pumped through the flow tube using a peristaltic pump at different flow rates, and the temperature of the fluid was maintained at 4°C. Actual experimental flow rates were determined by manual calibration of pump flow settings using a stopwatch to record the time required to deliver a measured fluid volume. Dean flow pattern in the curved tube critically influences the efficiency of the fluid mixing by the centripetal forces at the curve, which is governed by Dean number (*D_e_*).

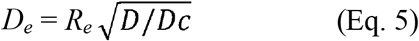

**Figure 1.**
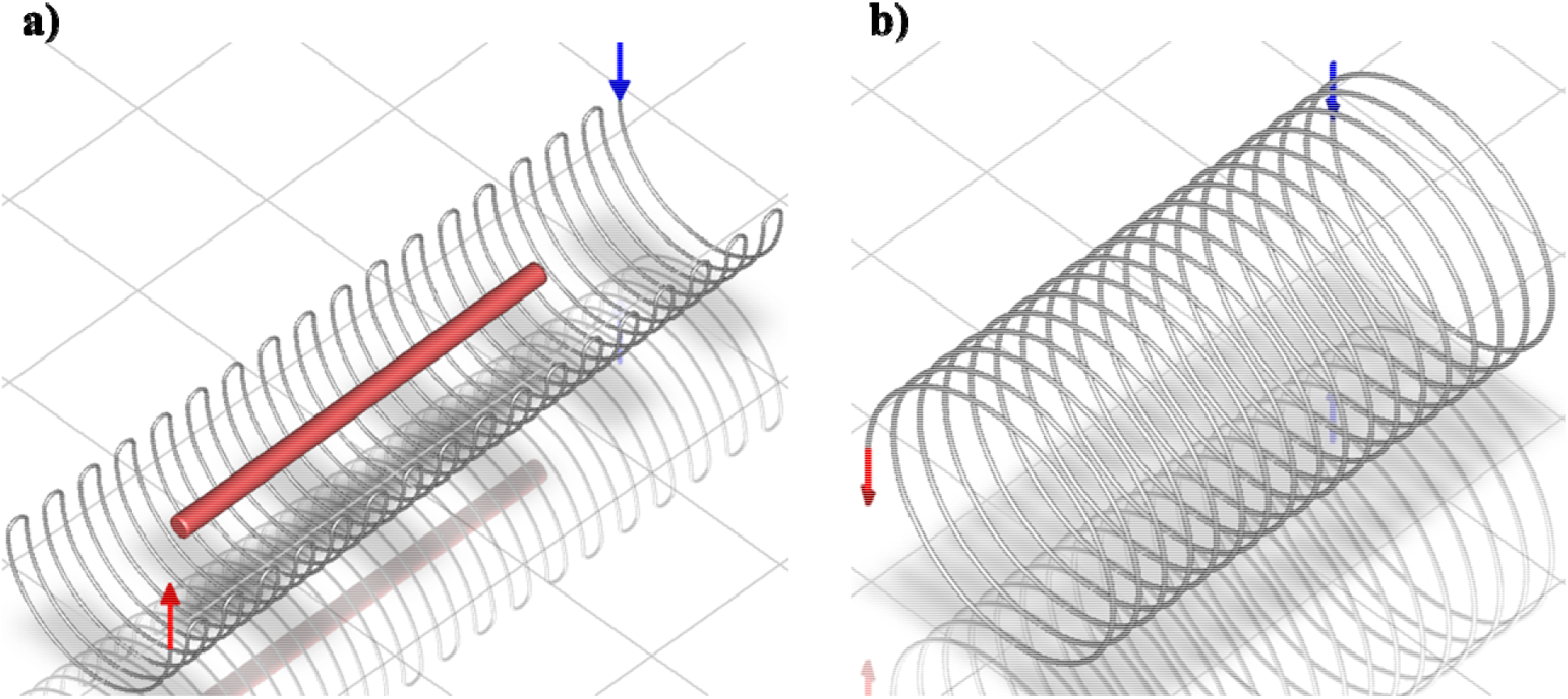
Geometric model of the UV-C reactors; a) SPCT; b) HPCT

Where *D* is the tube diameter (0.00159 m), *R_e_* represents the tube Reynolds number, and *D_c_* is the curvature diameter (0.01 m at the serpentine bend, 0.09 m at coil region).

### 2.6 Numerical simulation of fluid velocity magnitude and incident radiation

CFD software Ansys Fluent 2020 R2 (Ansys Inc., PA, United States) commercial version was used to compute velocity magnitude and incident radiation fields in serpentine and helical coil tube reactors. The 3D geometric model of the reactors with the same area and coil diameter was created using the Ansys Space Claim (Figure 1). Then the geometry was imported into the Ansys meshing tool, which discretizes the geometries into smaller finite tetrahedron-shaped elements (> 3.8 million) where the governing equations were resolved. Inflation was used to get finer mesh near the tube wall. The meshed geometry was imported into the Ansys FLUENT solver tool for numerical simulations. Based on Re, laminar flow conditions were assumed within the reactor, and a laminar viscous model was selected. The Continuity equation expresses the conserved quantity transport and Reynolds-averaged Navier Stokes equations for incompressible flows in equations 6 and 7.

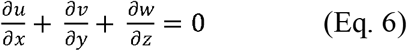

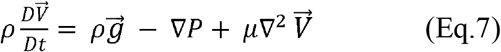

Where ∇*P* is the pressure gradient (Pa), *g* is the gravitational acceleration (m/s^2^).

The discrete ordinate (DO) model (Eq. 8) was applied for incident radiation simulations as it solves the radiation transfer equation over several discretized solid angles with certain directional vectors. The angular discretization of the model was set at 8×8 divisions and 6×6 pixelations, and the wavelength of the radiation band was chosen as 254 nm. Fluid domain properties (density, viscosity, absorption and scattering coefficients, refractive index) for WM and AM were given in Table 1. Inlet boundary was assigned as velocity inlet with velocity 0.59, 1.41m/s for WM and AM, respectively. The outlet section was applied as set as a pressure outlet. The SIMPLE algorithm method was used for the pressure–velocity coupling and second-order upwind scheme applied for the momentum convection and discrete ordinates.

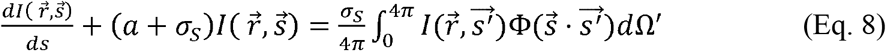

**Table 1:** Optical and physical properties of WM and AM *B. cereus* spore suspensions

| **Optical/physical properties** | **WM spore suspension** | **AM spore suspension** |
| --- | --- | --- |
| pH | 6.73 ± 0.03 | 6.8 ± 0.02 |
| Density (Kg/m<sup>3</sup>) | 1028 | 1140 |
| Viscosity (Pa·s) | 0.0019 | 0.012 |
| Absorption coefficient (base<sub>10</sub> 1/cm) | 24.4 ± 0.4 | 16 ± 0.3 |
| Scattering coefficient (base<sub>10</sub> 1/cm) | 44 ± 0.6 | 14.9 ± 0.2 |
| Reflectance (%) | 11.9 | 9.1 |
| Refractive Index | 1.4016 | 1.398 |

Where *I* is the radiation intensity, 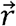 is the position vector, 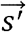 is the scattering direction vector, *s* is the path length, *a* is the absorption coefficient, *n* is the refractive index, *σ_s_* is the scattering coefficient, *σ* is the Stephen-Boltzmann constant, Φ is the phase function and Ω′ is the solid angle.

### 2.7 Analysis of lipid peroxidation

Malonaldehyde (MDA) (a usual end product of lipid peroxidation) concentrations in samples (control and UV exposed) were determined according to the method of Nielsen et al. (1997). WM or AM samples were suspended with 1% ortho-phosphoric acid and 42 mM thiobarbituric acid (TBA) using a vortex mixer, heated at 100 °C for 60 minutes reaction was terminated by ice-cooling for 10 min. Then 300 μL of the reaction mixture was transferred to microcentrifuge tubes containing 300 μl of 2 N NaOH: MeOH (1:12, v/v) solution and centrifuged for 3 min at 13,000 rpm. The supernatant was passed through a 0.2 μm Nylon membrane filter, and MDA concentration was read at 532 nm. 1,1,3,3-Tetraethoxypropane (TEP), which stoichiometrically releases MDA as end product upon acid hydrolysis, was used as a standard. Thiobarbituric acid reactive substances (TBARS) values of the samples were estimated from a standard curve and reported as mg MDA per kg test fluid (MDA/kg).

### 2.8 Analysis of volatile compounds

A Flash GC electronic nose (Heracles II from Alpha MOS, Toulouse, France) was used to analyze volatile aroma compounds (Pendyala et al., 2020). Untreated and UV-C treated watermelon beverage samples (10 mL) were placed in 20 mL vials and then sealed with an airtight septa screw cap using a crimper. The samples were equilibrated then headspace aroma generated in the vials was injected using an autosampler into the electronic nose column at a flow rate of 270 μL s^-1^. For calibration, an alkane solution (C6-C16) was used. The analysis was repeated three times, and data were analyzed by inbuilt AlphaMOS software. The individual volatile compounds, including Kovats index, retention time, sensory description, were identified using inbuilt software (AromaChemBase).

### 2.7 Statistics

A balanced design with three replicates for each treatment was exposed to the selected UV irradiation treatment. Each sample was independent and assigned randomly to a treatment. Oneway ANOVA with Tukey’s HSD multiple comparison tests was performed to assess the effects of different treatments on microbial inactivation, lipid peroxidation, and volatiles. Data are presented as means ± one standard deviation from the mean. Statistical significance was tested at a 5 percent significance level (p > 0.05).

## 3. Results and Discussion

### 3.1 Optical and physical properties of WM and AM spore suspensions

The optical and physical properties of WM and AM spore suspensions are shown in Table 1. The pH, viscosity, and density of the WM spore suspension were measured as 6.73 ± 0.03, 0.0019 Pa·s, and 1028 Kg/m^3^, respectively. Absorption, scattering coefficient, the reflectance of WM spore suspension at 254 nm was calculated as 24.4 ± 0.4/cm. 44 ± 0.6/cm and 11.9 %, respectively. For AM spore suspension, pH, viscosity, and density were found to be 6.8 ± 0.02, 0.012 Pa·s, and 1140 Kg/m^3^, respectively. The absorption, scattering coefficient, reflectance of AM test fluid at 254 nm was estimated to be 16 ± 0.3/cm. 14.9 ± 0.2, and 9.1 %, respectively.

The optical data indicate that WM was a strong absorber and scatterer of UV-C light compared to AM. Light scattering by fat globules and casein micelles causes milk to appear turbid and opaque. Light scattering occurs when the wavelength of light is close to the same magnitude as the particle. Thus, smaller particles scatter light of shorter wavelengths. These light attenuation properties of WM and AM affect the penetration depth of UV-C photons and require turbulent mixing conditions for efficient fluence distribution to microbial particles (Pendyala et al., 2021, Vashisht et al., 2021). Hence penetration depth and fluid mixing are crucial parameters for designing and developing an efficient reactor system for processing WM and AM.

### 3.2 Design of an SPCT UV-C system and comparative numerical simulation of the flow field and radiation field distribution in SPCT and HPCT reactors

Our previous studies demonstrated that efficient UV-C inactivation of microorganisms in beverages (coconut water, cranberry flavored water, diluted milk, and watermelon juice) with a UV-C 254 nm absorbance range of 1.1 – 9.8 AU/cm using a dean flow reactor system with tube diameter 3.175 mm (Bhullar et al., 2018; Ward et al., 2018; Gopisetty et al., 2018, 2019; Pendyala et al., 2020). In this investigation, by considering high absorbance, scattering coefficients, and reflectance of WM and AM, we selected a tube diameter of 1.59 mm with sharp serpentine bends (pitch 1 cm) for effective light penetration and fluid mixing using CFD. To compromise the residence time and fluid mixing, experimental flow rates of 70 and 162 mL/min for WM and AM, respectively, were selected. As the viscosity of the AM was > 6 folds higher than the AM, we selected a high flow rate (162 mL/min) for AM experiments. Re, De, and fluid velocity at selected flow rates is given in Table 2.

**Table 2:** Flow parameters of WM and AM *B. cereus* spore suspensions

| **Flow parameter** | **WM spore suspension** | **AM spore suspension** |
| --- | --- | --- |
| Flow rate (mL/min) | 70 | 162 |
| Reynolds number | 510 | 213 |
| Dean number at bend location | 203 | 85 |
| Dean number at other curved location | 68 | 29 |
| volume average Velocity magnitude (m/s) | 0.59 | 1.41 |

The WM flow field distribution inside the SPCT and HPCT reactors with the same area and pitch (1 cm) were simulated at an inlet flow rate of 70 mL/min. The velocity magnitude of both reactors at different locations was compared (Figure 2). The contour plots clearly show no significant difference in velocity magnitude distribution between SPCT and HPCT reactor outlet locations (Figure 2a & 2b). In contrast, efficient velocity distribution was observed at the serpentine bend geographical location of SPCT (Figure 3a & 3b). The calculated De at the serpentine bend location was 203, which is ~3 times higher than the other curve with De 68. At De > 64~75, the Dean vortices become stable, which confers a primary dynamic instability, a secondary instability occurs for De > 75~200, followed by the appearance of transition to turbulence with full turbulence for De > 400 (Ligrani, 1994).

**Figure 2.**
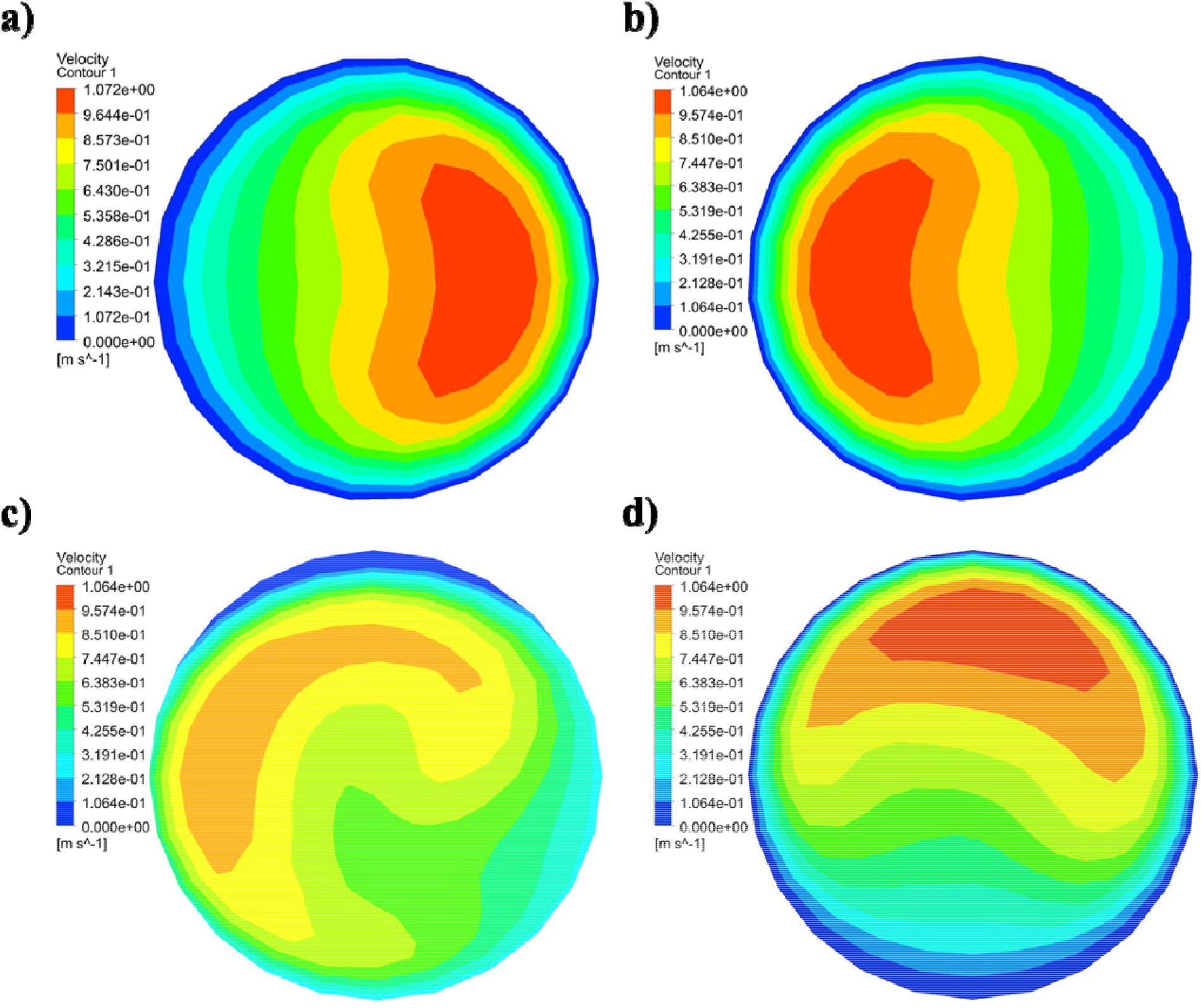
Velocity magnitude contours of WM UV-C reactors; a) HPCT reactor; b) SPCT reactor; c) SPCT reactor at 60 ° angle to serpentine bend end; d) SPCT reactor at serpentine bend end

**Figure 3.**
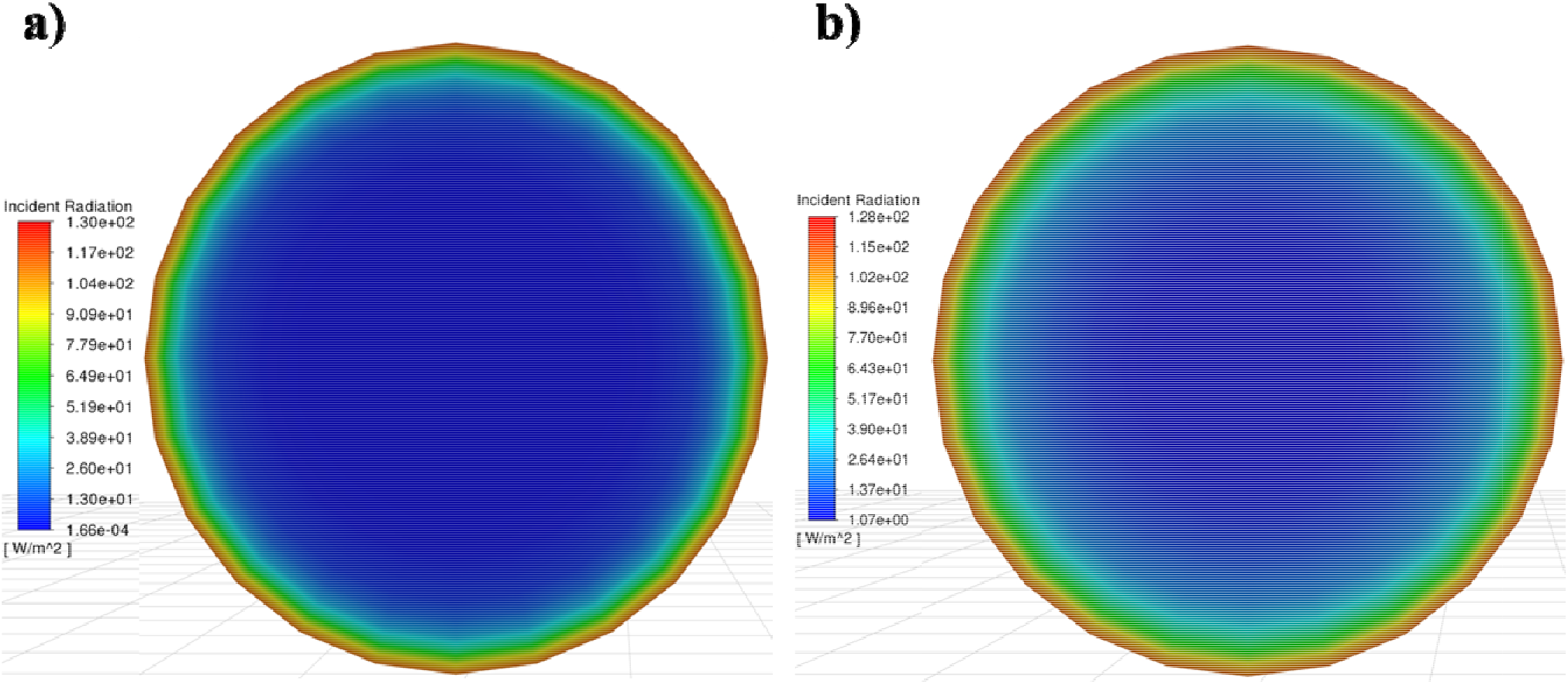
UV-C radiation contours of WM and AM; a) WM; b) AM

UV-C radiation distribution in WM and AM is shown in Figure 3. The radiation intensity gradually decreases from the wall towards the center of the tube due to light attenuation by WM or AM. As UV-C absorption and scattering of the WM were noticed to be higher than AM (Table 1), the radiation-free dead space in the reactor was higher with WM compared to the AM. So, it is essential to mix radiation-free zone with radiation zone to provide efficient UV-C fluence distribution throughout the reactor.

### 3.3 Dose-response inactivation kinetics of *B.cereus* spores inactivation in WM and AM

The scattering factor calculated for WM (12 %) was higher than the AM (7.2 %) and considered for delivered UV dose calculation. Figure 3 shows the UV dose-response curves of *B. cereus* spores inactivation in WM and AM obtained with a standard collimated beam apparatus (USEPA, 2006). The data was best fitted to a polynomial quadratic regression equation with the goodness of fit (*R^2^* > 0.99). The UV sensitivity (*D*_10_) of *B. cereus* spores in WM and AM was estimated as 8.8 and 9.1 mJ/cm^2^, respectively. These data were well correlated with the literature studies reported values (8.75 – 9.29 mJ/cm^2^) in phosphate buffer saline and coconut water (Blatchley et al. 2005, Pendyala et al., 2019 & 2020). The inactivation model coefficients from dose-response equations (Figure 3) were used to estimate REF delivered by SPCT UV system by substitution of ‘x’ with log reduction results. Literature studies reported delivered UV fluence in two ways, i.e., average fluence and REF. However, the average fluence calculated based on only optical attenuation properties of fluids without fluid flow field is not suitable for continuous flow conditions. The fluid flow field is the crucial parameter that can influence delivered fluence. Therefore, the term REF, which relies on microbial inactivation under experimental conditions, was used in this study.

### 3.3 UV-C inactivation of *B. cereus* spore suspensions of WM and AM

WM spore suspension was treated with the SPCT UV-C system for six passes, and samples were sampled at passes 1, 3, 5, 6. Since the viscosity of AM was higher than WM, the AM spore suspension was passed through the UV-C system from 0 to 10 passes and sampled at pass 2, 4, 6, 10. Figure 4a shows the inactivation of *B. cereus* endospores suspended in WM. The data revealed that 1.1 ± 0.27, 2.7 ± 0.15, 3.4 ± 0.10, and 4.2 ± 0.05 log CFU/mL reduction of *B. cereus* endospores in WM was observed and thereby delivered 7.7 ± 1.86, 21 ± 1.89, 27 ± 0.85, and 33 ± 0.73 mJ/cm^2^ of REF at passes 1, 3, 5 and 6, respectively. Inactivation data of *B. cereus* endospores suspended in AM is given in Figure 4b. Experimental data show at passes of 2, 4, 6 and 10, *B. cereus* endospores in AM reduced to 0.74 ± 0.1, 1.8 ± 0.12, 2.5 ± 0.24, and 4.0 ± 0.12 log CFU/mL, and delivered 6.8 ± 0.89, 16.4 ± 1.14, 22.6 ± 2.47, and 36.5 ± 1.9 mJ/cm^2^ of REF, respectively. The results revealed that ≥ 4 log reduction of *B. cereus* endospores was noticed after six passes in WM at 70 mL/min flowrate, whereas the same was noticed after ten passes in AM at 162 mL/min. In our earlier work, > 6 log reduction of *B. cereus* endospores was reported at UV-C REF of 60 mJ/cm^2^ in 50 % watermelon beverage treated with continuous dean flow system (Pendyala et al. 2020). Choudhary et al. (2011) and Bandla et al. (2012) reported 2.65 (in raw cow milk), 2.72 (in skim cow milk), and 3.29 log reduction (in raw soy milk) of *B. cereus* spores by treatment with coiled tube 1.6 mm ID UV system. But the authors did not measure optical light attenuation parameters and not validate the delivered UV fluence under experimental conditions.

**Figure 4.**
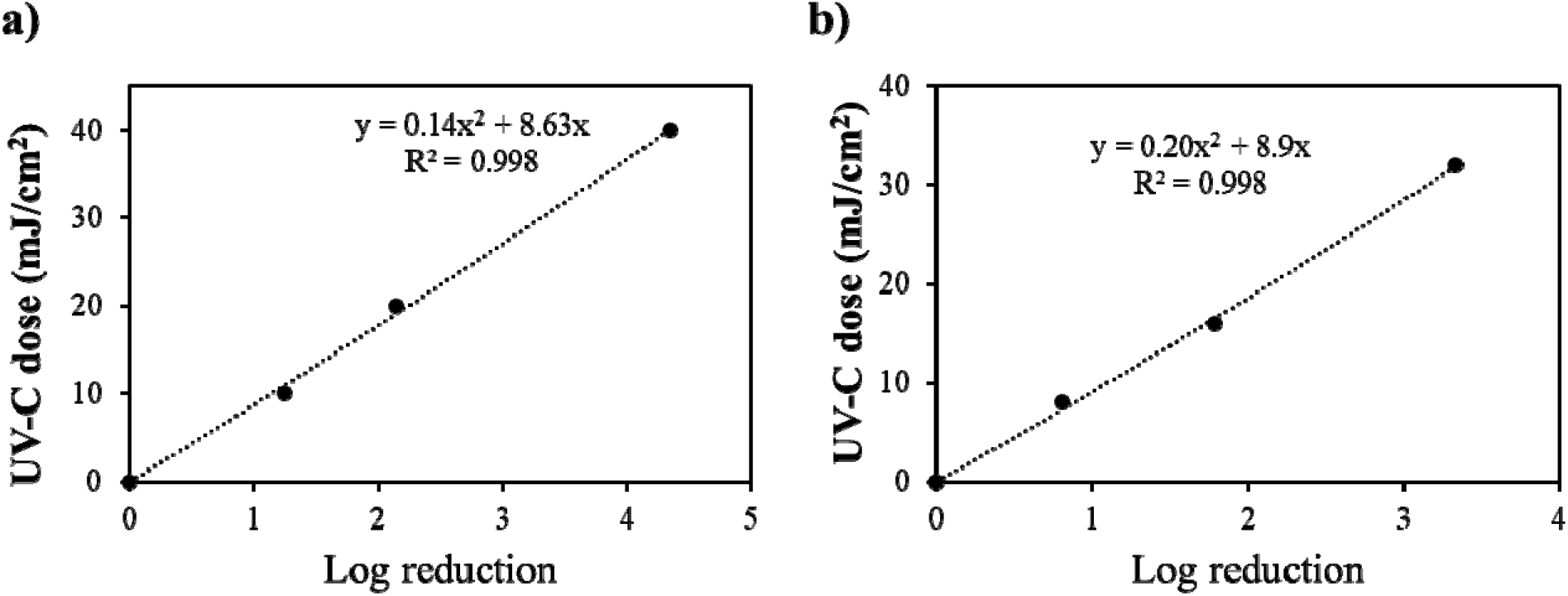
Dose response curves of B. *cereus* endospores suspended in WM and AM; a) WM; b) AM

**Figure 5.**
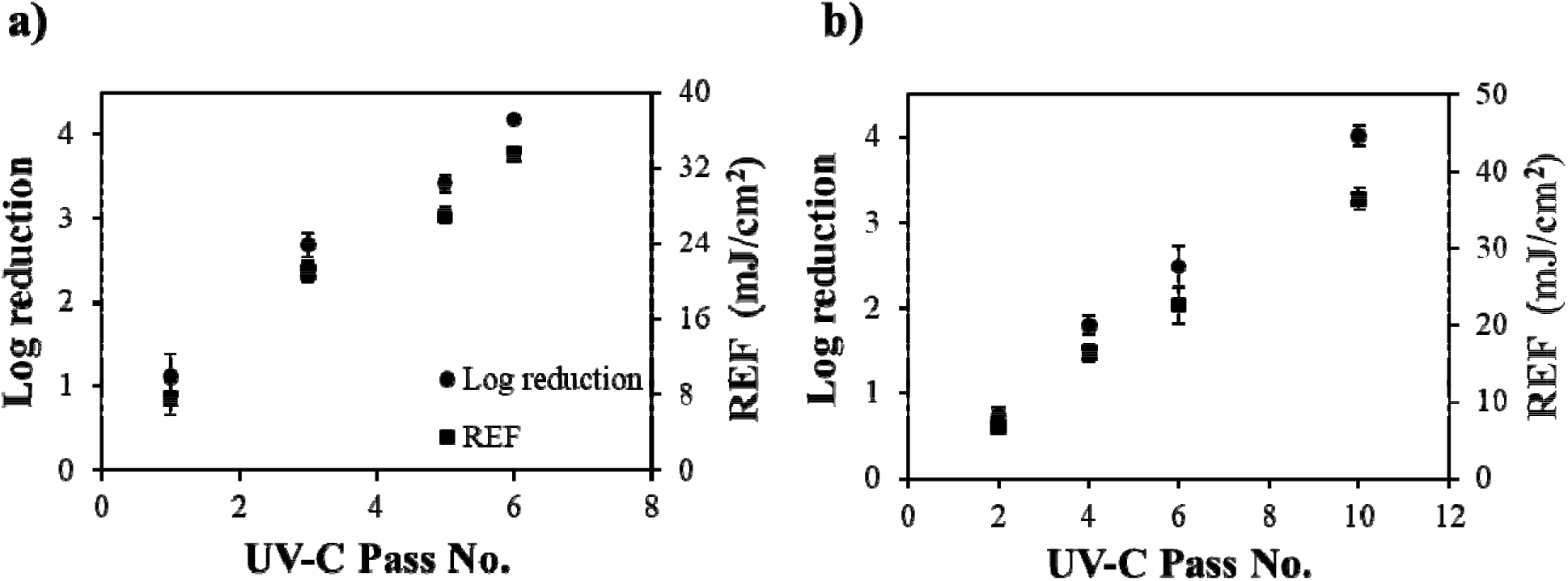
Inactivation data of B. *cereus* endospores with designed SPCT UV-C reactor; a) WM suspension; b) AM suspension

### 3.4 Assessment of UV-C treated WM and AM quality

#### 3.4.1 Lipid peroxidation

Milk contains numerous chemical compounds that can involve in photoreactions. During exposure to UV light, photooxidation of milk occur either by direct photolysis (by the production of free radicals primarily from lipids) or by photosensitized oxidation (occurs in the presence of photosensitizers) (Bradley and Min, 1992). However, direct photolysis is minimal and, thus, not a major concern (Frankel, 2005). Photosensitizers can absorb UV or visible light to become electronically excited states: singlet and triplet. The life span of the triplet excited state is longer and initiates oxidation. Photosensitizer mediated photooxidation can happen through either type I (through a free radical mechanism) or type II (reacts with oxygen to produce highly reactive singlet oxygen) reactions (Frankel, 2005). Riboflavin, protoporphyrin IX (PpIX), hematoporphyrin, a chlorophyll a-like compound, and two unidentified tetrapyrroles are photosensitizers present in the milk (Wold et al., 2005). Among these riboflavins that absorb UV light with absorption maxima at 266 nm is the potent UV-C photosensitizer in milk. Hence evaluation of lipid peroxidation is the crucial parameter to determine the quality of UV-C irradiated milk. The TBARS values of control WM was noticed to be 0.28 ± 0.04 mg MDA/kg whereas UV irradiated WM at 60 mJ/cm^2^ shown slight increase 0.43 ± 0.07 mg MDA/kg. These results are in the range of data reported for goat milk (0.31 – 0.58 mg MDA/kg) and cow milk (0.59 – 0.92 mg MDA/kg) (Matak et al., 2007, van Aardt et al., 2005). However, the delivered fluence was not verified in both studies.

#### 3.4.2 Volatile compounds profile

Control and UV irradiated samples were maintained at 4 °C until volatiles analysis. The volatile compounds profile in WM and AM samples was analyzed using an electronic nose by comparing their retention times with Kovats retention indices (Śliwińska et al., 2016). The volatile compounds profile of UV-C treated WM is shown in Table 2. The results revealed that esters, aldehydes, and alcohols accounted for >75 % of the volatile compounds. 1-Propanol, 2-methyl propanal, butanal, and ethyl 2-methyl butyrate were identified as major volatile compounds in untreated WM. Compared to untreated samples, as UV fluence increased, the proportion of aldehydes, alcohols, and ethers decreased in total volatiles. In contrast, a relative increase in other compounds such as esters, ketones, and acids was observed. Comparatively within UV-C treated samples (16 – 64 mJ/cm^2^) revealed that the proportion of esters, aldehydes, alcohols, and ketones in total volatiles were slightly altered. Table 3 shows the volatiles profile of AM and UV-C treated AM. Methyl-2-methyl butanoate, Benzaldehyde, 2-Methylbutane, Ethyl 2-methyl butyrate, and Butanal were occupied > 56 % in total volatiles. In response to UV-C radiation, no significant change (p > 0.05) in the total proportion of aldehydes, alcohols, esters, ketones, and others compounds was observed. However, benzaldehyde proportion in the aldehyde decreased whereas the hexanal proportion increased in the UV-C treated samples. Further investigation may unveil the mechanism for the increase in hexanal and decrease in the benzaldehyde. Overall, the volatile compounds of irradiated AM and WM were well retained.

**Table 3:**
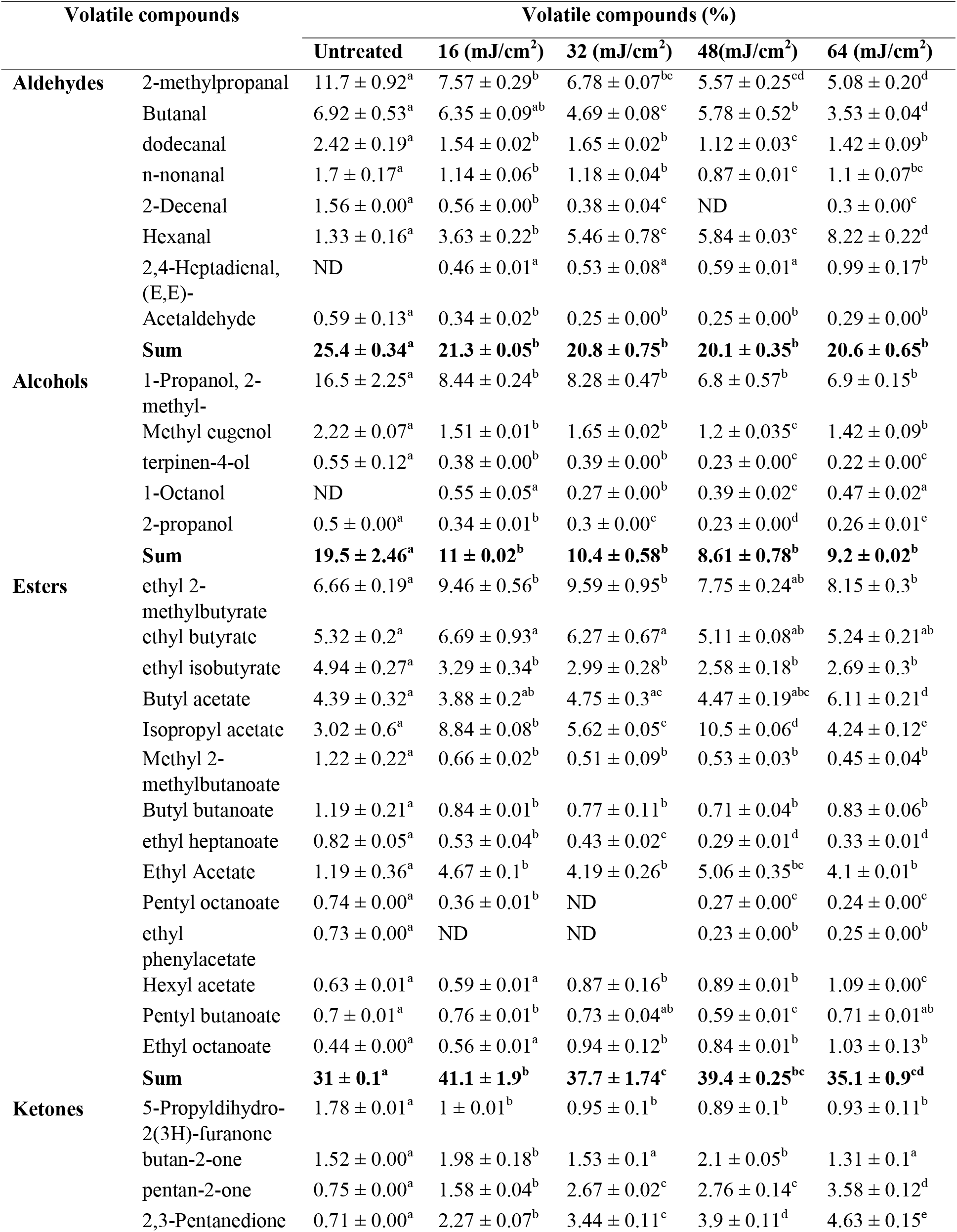
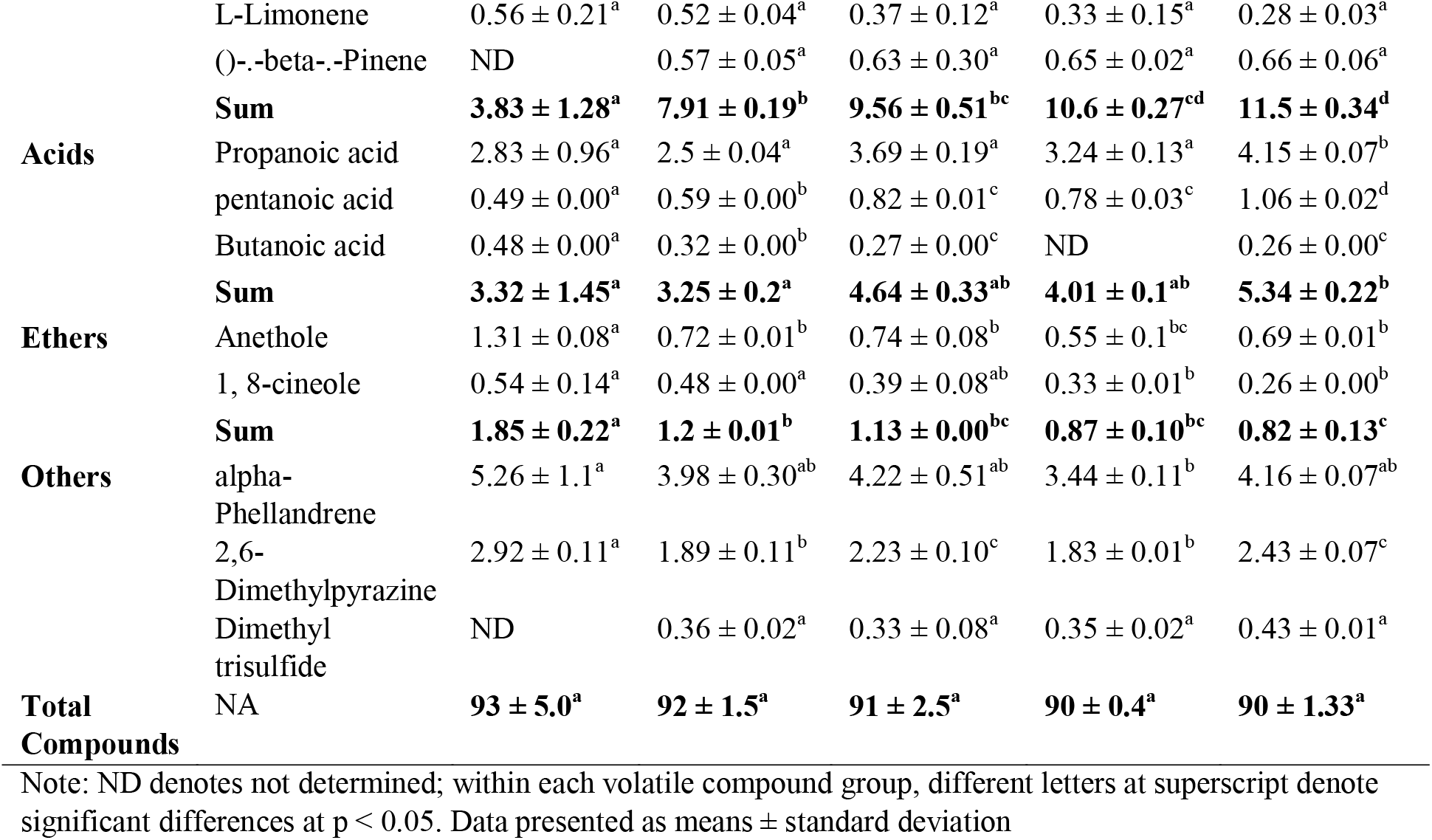
GC-Surface percentage of volatile compounds in UV-C treated whole cow milk.

**Table 4:** GC-Surface percentage of volatile compounds in UV-C treated Almond milk.

| Volatile compounds |  | Volatile compounds (%) |  |
| --- | --- | --- | --- |
|  |  | Untreated | 60 mJ/cm<sup>2</sup> |
| **Aldehydes** | Benzaldehyde | 10.41 ± 0.81<sup>a</sup> | 7.47 ± 0.58<sup>b</sup> |
|  | Butanal | 4.76 ± 0.23<sup>a</sup> | 3.86 ± 0.2<sup>b</sup> |
|  | 2-methyl propanal | 3.8 ± 0.18<sup>a</sup> | 2.9 ± 0.15<sup>b</sup> |
|  | Dodecanal | 1.85 ± 0.11<sup>a</sup> | 1.25 ± 0.1<sup>b</sup> |
|  | Hexanal | 3.24 ± 0.23<sup>a</sup> | 9.83 ± 0.77<sup>b</sup> |
|  | 3-methyl butanal | 3.68 ± 0.21<sup>a</sup> | 4.55 ± 0.27<sup>b</sup> |
|  | 2,4-Heptadienal, (E,E)- | 0.9 ± 0.03<sup>a</sup> | 1 ± 0.02<sup>b</sup> |
|  | n-nonanal | 0.92 ± 0.02<sup>a</sup> | 0.7 ± 0.02<sup>b</sup> |
|  | Propanal | 0.78 ± 0.02<sup>a</sup> | 0.76 ± 0.02<sup>a</sup> |
|  | **Sum** | **30.34 ± 1.84<sup>a</sup>** | **32.34 ± 2.13<sup>a</sup>** |
| **Alcohols** | 2-propanol | 4.69 ± 0.25<sup>a</sup> | 4.52 ± 0.29<sup>a</sup> |
|  | Methyl eugenol | 3.35 ± 0.21<sup>a</sup> | 2.56 ± 0.19<sup>b</sup> |
|  | **Sum** | **8.04 ± 0.46<sup>a</sup>** | **7.08 ± 0.48<sup>a</sup>** |
| **Esters** | Methyl-2-methylbutanoate | 27.47 ± 1.85<sup>a</sup> | 24.81 ± 1.92<sup>a</sup> |
|  | Ethyl 2-methylbutyrate | 6.33 ± 0.43<sup>a</sup> | 5.69 ± 0.51<sup>a</sup> |
|  | Ethyl isobutyrate | 2.63 ± 0.17<sup>a</sup> | 1.19 ± 0.11<sup>b</sup> |
|  | Butanoic acid | 0.95 ± 0.1<sup>a</sup> | 2.16 ± 0.16<sup>b</sup> |
|  | Isoamyl acetate | 1.96 ± 0.13<sup>a</sup> | 1.63 ± 0.18<sup>a</sup> |
|  | **Sum** | **39.34 ± 2.68<sup>a</sup>** | **35.48 ± 2.88<sup>a</sup>** |
| **Ketones** | 2,3-Pentanedione | 1.37 ± 0.09<sup>a</sup> | 1.28 ± 0.1<sup>a</sup> |
| **Carboxylic Acid** | 2-methylpropanoic acid | 3.5 ± 0.27<sup>a</sup> | 3.13 ± 0.29<sup>a</sup> |
|  | 3-methylbutanoic acid | 1.45 ± 0.11<sup>a</sup> | 1.28 ± 0.17<sup>a</sup> |
|  | **Sum** | **4.95 ± 0.38<sup>a</sup>** | **4.41 ± 0.46<sup>a</sup>** |
| **Saturated hydrocarbon** | 2-Methylbutane | 7.43 ± 0.42<sup>a</sup> | 6.22 ± 0.47<sup>a</sup> |
| **Terpenes** | 1R-(+)-alpha-pinene | 1.52 ± 0.12<sup>a</sup> | 1.47 ± 0.15<sup>a</sup> |
|  | ()-.beta-.Pinene | 0.8 ± 0.07<sup>a</sup> | 0.76 ± 0.1<sup>a</sup> |
|  | Thymol | 1.32 ± 0.1<sup>a</sup> | 0.78 ± 0.06<sup>b</sup> |
|  | **Sum** | **3.64 ± 0.29<sup>a</sup>** | **3.01 ± 0.31<sup>a</sup>** |
| **Ethers** | Dibenzyl ether | 1.02 ± 0.1<sup>a</sup> | 0.86 ± 0.1<sup>a</sup> |
| **Total compounds** | **NA** | **96.1 ± 6.3<sup>a</sup>** | **90.68 ± 6.91<sup>a</sup>** |

Note: ND denotes not determined; within each volatile compound group, different letters at superscript denote significant differences at $p < 0.05$ . Data presented as means ± standard deviation

## 4. Conclusions

This study designed and developed a novel SPCT UV system and demonstrated effective inactivation (> 4 log CFU/mL) of endospores in WM and AM using a thin-film serpentine coiled tube UV-C reactor System. TBARS values and volatiles profile were not altered significantly, indicating that light-sensitive lipid compounds and light and temperature-sensitive volatile compounds were well retained in the UV irradiated WM and AM. Further studies on the design of SPCT UV system with adequate flow rate and serpentine bends to provide turbulence (De ~ 400) in the reactor and subsequent development of a scale-up model to process fluids at higher flow-rates will be warranted in the future.

## CRediT authorship contribution statement

**Brahmaiah Pendyala:** Conceptualization, Methodology, Investigation, Visualization, Writing – original draft. **Ankit Patras:** Conceptualization, Methodology, Supervision, original draft review, Funding acquisition. **Vybhav Vipul Sudhir Gopisetty**: Methodology, Investigation. **Pranav Vashisht:** Methodology, Investigation. **Ramasamy Ravi:** Methodology, Investigation. Note: There are no conflicts to declare.

## Acknowledgments

This project was funded under the Agriculture and Food Research Initiative (Food Safety Challenge Area), USDA, Award numbers; 2018-38821-27732, and 2019-69015-29233. The authors would like to thank Trojan Technologies for providing valuable guidance in this project.

